# Mechanical constraint causes lower turgor, thicker walls, and faster growth in Arabidopsis root hairs

**DOI:** 10.1101/2025.11.25.690544

**Authors:** Vyankatesh Zambare, Hojae Yi, Charles T. Anderson

## Abstract

Root hairs absorb water and mineral nutrients while anchoring growing root tips. They must navigate through soils of varying mechanical properties. Mechanical changes in their microenvironment affect root hair shape and growth rate, but what underlies these responses has remained elusive. To uncover these mechanisms, we grew seedlings of *Arabidopsis thaliana* (Col-0) in media with increasing mechanical stiffness. Col-0 seedlings showed both shorter and fewer root hairs with increasing mechanical resistance. We used incipient plasmolysis to estimate turgor pressure in trichoblasts and found that it also decreased with increasing media stiffness. Because cellulose is the major load-bearing polymer in the cell wall and influences cell expansion, we quantified cellulose orientation in root hairs. We found that cellulose fibrils were oriented at steeper angles relative to the growth axis in root hairs grown in stiffer media, suggesting a response to mechanical stress. Cell wall thickness also increased with increasing media stiffness. Microtubule orientations followed patterns that were similar to those of cellulose fibrils, but at a smaller angle relative to the growth axis, whereas microtubule density decreased with increasing media stiffness. Unexpectedly, we observed that root hairs grew faster in stiffer media, implying that their growth is misregulated, potentially triggering wall integrity signaling that causes early growth arrest. Finite element modeling of root hairs predicted decreased surface stress, explaining these growth phenotypes. These findings help establish mechanistic links between mechanotransduction, cytoskeletal dynamics, and cell wall assembly during root hair growth.

## Introduction

Root hairs are tubular projections of epidermal cells on the root surface. They are primarily found in the maturation zone of roots and increase root surface area, enhancing the absorption of water and mineral nutrients (Grierson et al., 2014; Grierson & Schiefelbein, 2002). Root hairs also help anchor the growth of root tips and assist in navigating through soil pores and penetrating compact soils (Bengough et al., 2016). In response to chemo-mechanical cues, root hairs can undergo dynamic changes in shape and growth (Pereira et al., 2024). Root hairs also exhibit variable phenotypes in response to soil texture and elemental composition (Ma et al., 2001; Zhou et al., 2024). Therefore, root hairs are beneficial for plant survival and display morphogenetic plasticity.

Root hair development consists of two main stages – cell fate determination and morphogenesis – and is regulated by a well-characterized genetic network (Balcerowicz et al., 2015). Epidermal cells that form root hairs are called trichoblasts, whereas those that lack root hairs are called atrichoblasts. WEREWOLF, a MYB transcription factor, is preferentially expressed in atrichoblasts and is a positive regulator of GLABRA2. In the adjacent epidermal cells, trichoblasts, CAPRICE, a MYB-like protein, inhibits WERWEREWOLF expression. The interaction between WERWEREWOLF, CAPRICE, and GLABRA2 creates feedback loops that help maintain the established cell fate pattern (Schiefelbein et al., 2014). In many plant species, including the model plant *Arabidopsis thaliana*, trichoblasts develop in the cleft between two underlying cortical cells, whereas atrichoblasts are located over a single cortical cell (Lee & Schiefelbein, 1999). This patterning implies that mechanical and/or signaling interactions between the cortex and epidermis might help determine trichoblast fate, but the mechanisms underlying this patterning are not fully uncovered.

After trichoblast identity is established, root hairs initiate and grow from the rootward ends of these cells. This involves the reorganization of cellular and secretory machinery within the trichoblast to build the growing tube of the root hair (Denninger et al., 2019; Vissenberg et al., 2001, 2020). Root hairs possess cytoskeletal networks extending from the trichoblast body along the shank to the tip of the root hair tube. Actin filaments are major elements of the root hair cytoskeleton and are organized into thick bundles that run parallel to the root hair shank. Actin-myosin dynamics also drive an inverse-fountain-like flow of cytoplasmic streaming (Ketelaar et al., 2003; Miller et al., 1999). Actin thus serves as a primary conduit for polarized membrane trafficking in root hairs. Vesicles containing cell wall material are delivered to the root hair tip via the actin cytoskeleton, enabling rapid growth (Szymanski & Cosgrove, 2009). Microtubules, on the other hand, are required to maintain cell polarity and directed tip growth in root hairs (Bibikova et al., 1999). Cortical microtubules are oriented parallel to the growth axis of the root hair, whereas endoplasmic microtubules run from the perinuclear cytoplasm toward the tip. Microtubule orientation dynamically changes throughout root hair growth and stabilizes in mature root hairs (Van Bruaene et al., 2004). Depolymerizing microtubules slows, then halts root hair growth, but allowing microtubule re-polymerization results in the resumption of growth (Bibikova et al., 1999; Brueggeman et al., 2022; Dupouy et al., 2025). Therefore, cytoskeletal networks play indispensable roles in root hair growth.

The cell walls of root hairs are also major determinants of their growth dynamics and shape. Cell wall thickness helps determine the mechanical properties of root hairs and influences their growth and physiology (Baez et al., 2025; Galway, 2006; Pereira et al., 2023); thicker cell walls prevent cell bursting but also impede the influx of water and minerals (Delmer et al., 2024). Like those of other growing cells, the walls of root hairs are composed of cellulose, hemicelluloses, and pectins, each of which makes unique contributions to cell wall mechanics (Cosgrove, 2022). Cellulose, a β-1,4-linked polymer of glucose that exists in multi-chain microfibrils in the plant cell wall, has high tensile strength and restricts cell expansion perpendicularly to its major axis of deposition (McNamara et al., 2015). In root hairs, cellulose is produced by both CELLULOSE SYNTHASE and CELLULOSE SYNTHASE-LIKE D glycosyltransferases, and COBRA-LIKE proteins are also involved in its production, especially at the root hair tip (Bernal et al., 2008; Galway et al., 2011; Li et al., 2022; Park et al., 2011; Peng et al., 2019). Hemicelluloses, which have equatorially linked sugar backbones and can contain side chains, are critical for root hair growth, especially in the case of xyloglucan, since mutants lacking this hemicellulose show short, bulged root hairs (Cavalier et al., 2008; Eckardt, 2008; Peña et al., 2012; Zabotina et al., 2012). Pectins, which are acidic polysaccharides, provide hydration and tunable flexibility to the wall (Cosgrove, 2022; Du et al., 2020). The anisotropic deposition of these wall polymers, with cellulose being laid down along linear trajectories by CELLULOSE SYNTHASES, which move along microtubule tracks, and hemicelluloses and pectins being secreted at the tip of the root hair, is hypothesized to result in a wall structure that resists radial expansion but facilitates axial growth, both at the root hair tip and along some regions of the root hair shank (Chen et al., 2016; Dumais et al., 2004; Herburger et al., 2022; Schoenaers et al., 2024). However, we do not fully understand how cell wall architecture is established in growing root hairs, or how this architecture relates to the mechanobiology of root hair growth and response to the physical environment.

Cell wall integrity has been implicated in the regulation of root hair growth; for example, the receptor-like kinase FERONIA, which binds to pectins and RAPID ALKALINIZATION FACTOR peptides, functions in root hair growth (Feng et al., 2018; Schoenaers et al., 2024). Additionally, recent studies of root hair mechanobiology using microfluidics have reported that root hair growth diminishes under greater mechanical resistance. In these experiments, the total growth duration is not affected, but the speed of growth decreases (Pereira et al., 2024). However, the reasons underlying these phenotypes are still unclear. This calls for exploring which changes in the subcellular components that control root hair growth might be responsible for growth responses in the face of mechanical resistance.

Here, we applied mechanically differing growth environments to study their effects on root hair morphogenesis. We found that roots tended to be shorter in stiffer growth media and formed shorter and fewer root hairs, resulting in decreased root hair density. Further analysis revealed a shift in epidermal cell patterning in response to increasing media stiffness that led to less trichoblast formation. Additionally, lower turgor pressure was observed in trichoblasts grown under increasing mechanical resistance. Under increasing media stiffness, cellulose in root hairs was oriented at a steeper angle relative to the growth axis, and cell wall thickness increased. Microtubule orientation followed a similar pattern to cellulose, and microtubule density decreased. In our conditions, the speed of root hair growth accelerated with increasing mechanical resistance, implying that the duration of growth is shorter in stiffer media. We combined our data into quantitative models of root hair growth that account for cell shape, wall mechanics, and turgor pressure. These models predict that wall stress is majorly modulated by turgor pressure and cell wall thickness, as root hair walls were under the lowest mechanical stress in the stiffest media. Together, these results help reveal the mechanobiological mechanisms by which root hairs navigate and respond to their physical environments.

## Results

### Increasing stiffness in the growth environment decreases root hair length and density

To study root hair morphogenesis under varying mechanical environments, we imaged roots of *Arabidopsis thaliana* (Col-0) seedlings grown into ½ MS media with differing agar concentrations. We observed that Col-0 roots produced shorter root hairs as the percentage of agar and thus the stiffness of the growth environment increased **(Fig. 1A-C)**. Fully elongated root hairs grown in 0.5% agar had an average length of 231.5 ± 49.71 µm (mean ± standard deviation), and those grown in 1% and 2% agar had average lengths of 161.7 ± 54.08 µm, and 111.3 ± 45.51 µm, respectively **(Fig. 1D)**. We also quantified lengths of epidermal cells and observed no significant change with increasing agar concentrations. Although, trichoblasts were always shorter than atrichoblasts **(Supp. Fig. S1)**. Col-0 seedlings also produced fewer root hairs per seedling with increasing agar content **(Fig. 1E)**: roots in 0.5% agar formed on an average 584.7 ± 114.1 root hairs per seedling, whereas those in 1% and 2% agar formed 314.4 ± 161.3 and 80.44 ± 46.41 root hairs, respectively. Decreased root hair number does not necessarily reflect lower root hair density, because external mechanical stress can also inhibit the elongation of the whole root. Therefore, we quantified root length and found that 5-day-old Col-0 seedlings produced shorter roots when grown in media with increasing agar content **(Fig. 1F)**. Roots grown in 0.5% agar were 13.74 ± 2.32 mm long on average, whereas those grown in 1% and 2% agar were 12.69 ± 2.97 mm and 10.85 ± 2.52 mm long, respectively. Due to the observed differences in root lengths, we then quantified root hair density expressed as the number of root hairs per mm of root. This value also decreased with increasing agar concentration **(Fig. 1G)**: roots grown in 0.5% agar had 42.8 ± 5.91 root hairs per mm, whereas roots grown in 1% and 2% agar had root hair densities of 24.92 ± 13.46 and 8.26 ± 7.09, respectively. Together, these data indicate that increasing media stiffness inhibits both root hair and root elongation.

**Fig 1.**
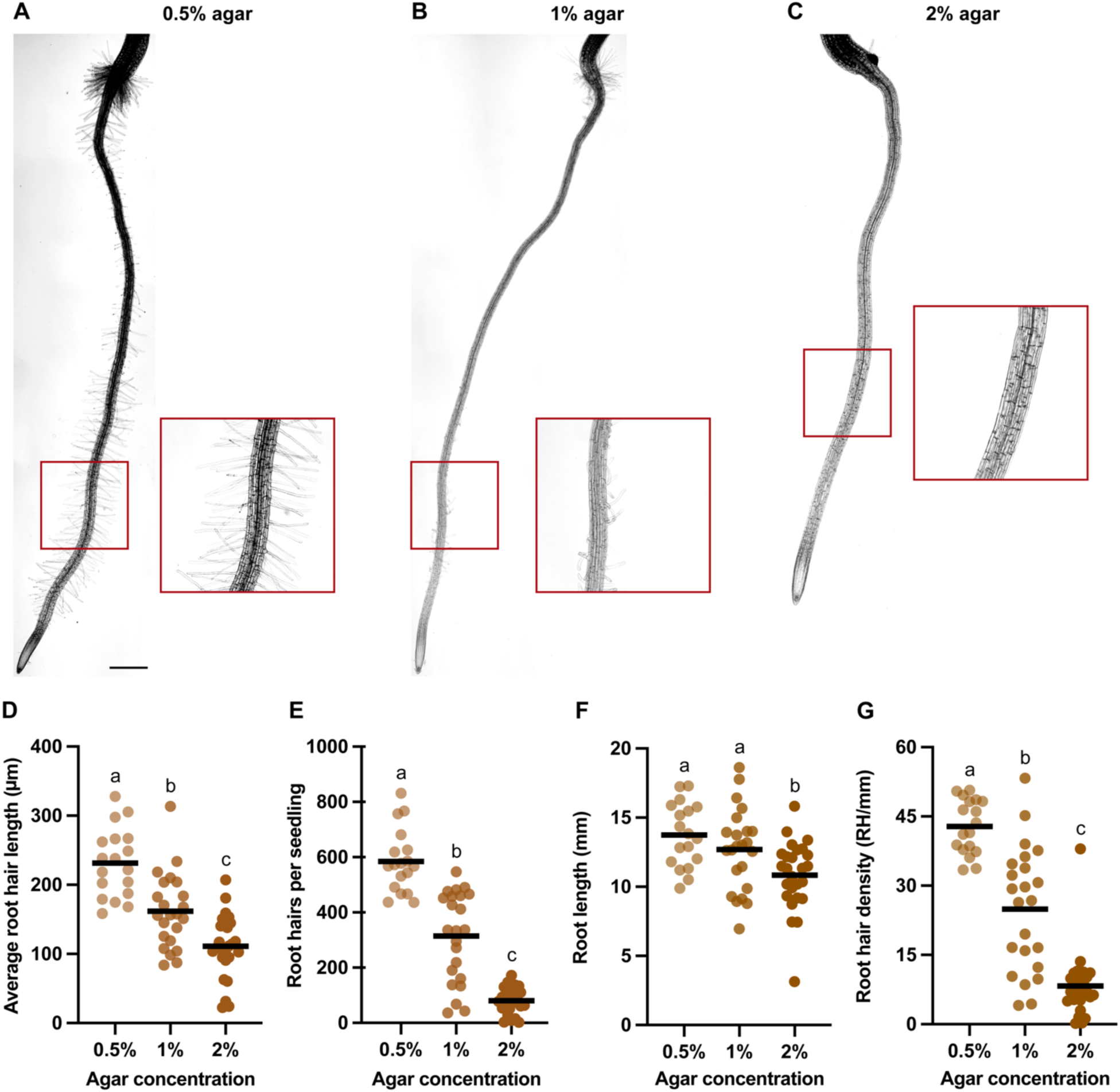
Root hair and root growth are inhibited by increasing stiffness of the growth media. **A-C,** Representative images of 5-day-old Col-0 roots grown in plant media supplemented with different concentrations of agar (Scale bar = 500 µm). **D,** Average root hair lengths, at least five root hairs measured per seedling. **E,** Root hair count per seedling. **F,** Whole root length. **G,** Root hair density represented by the number of root hairs per unit root length in mm. Each point in every graph represents one seedling. n = 18-26 seedlings per treatment from 5 independent experiments. Different letters denote significance, p < 0.05, ANOVA and post-hoc Tukey test.

### External mechanical stress perturbs epidermal cell patterning in roots by decreasing the proportion of trichoblasts

To explore the etiology of the root hair phenotypes we observed, we further quantified the developmental, biomechanical, cytoskeletal, and cell wall properties of Col-0 root hairs grown in mechanically different environments. Morphologically, trichoblasts are positioned at the junction between two cortical cells and contact both cortical cells (Dolan et al., 1994). Atrichoblasts, on the other hand, contact only one cortical cell at their inner periclinal surface **(Fig. 2A)**. Following these anatomical rules, we quantified the number of trichoblasts and atrichoblasts in 5-day-old Col-0 roots grown into media with increasing stiffness and found that the proportion of epidermal cells that are trichoblasts decreased with increasing agar concentration **(Fig. 2B)**. On average, 43.7% of the epidermal cells of roots grown in 0.5% agar were trichoblasts, a significantly higher percentage than in roots grown in 1% and 2% agar, which had 38.2% and 37.5% trichoblasts, respectively **(Fig. 2C)**. The proportion of atrichoblasts concomitantly increased with mechanical stress **(Fig. 2C),** but the number of epidermal cells and cortical cells remained constant across different agar treatments **(Supp. Fig. S2).** These data suggest that increasing mechanical stress induces a shift in epidermal cell patterning, resulting in fewer trichoblasts and helping to explain the observed decrease in root hair density **(Fig. 1G).**

**Fig 2.**
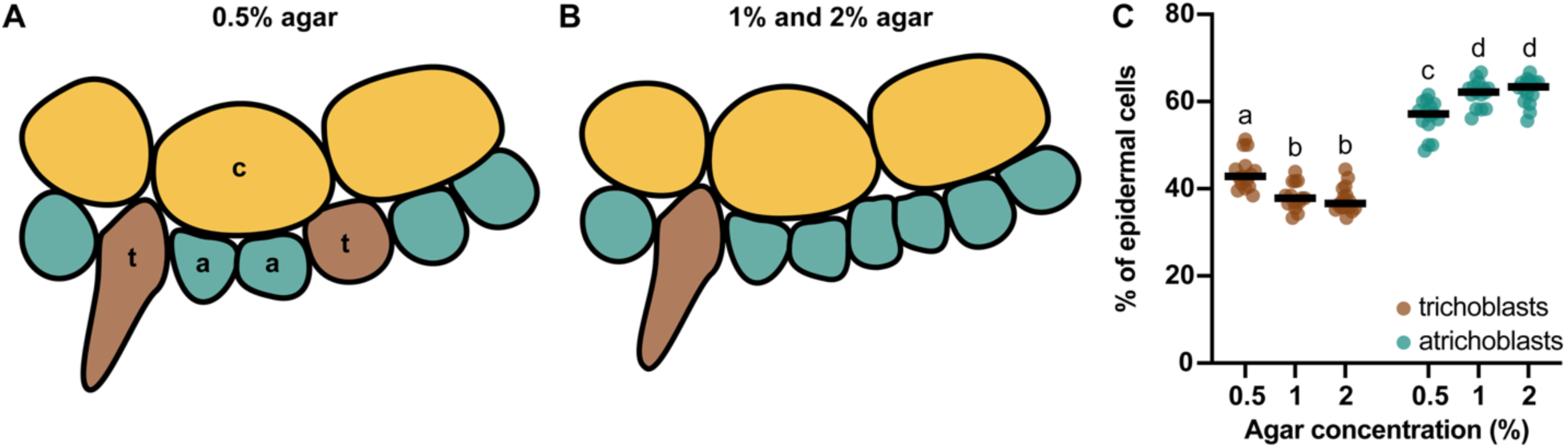
Increasing media stiffness perturbs cell patterning in the root epidermis. **A-B,** Illustration of epidermal cell pattering in transverse cross-sections of Arabidopsis roots grown in the softest and stiffest media tested. c-cortical cell, t-trichoblast, a-atrichoblast. **C,** The percentage of trichoblasts decreases and atrichoblasts increases with media stiffness. n ≥ 15 seedlings per treatment from 6 independent experiments. Different letters denote significance, p < 0.05, ANOVA and post-hoc Tukey test.

### Incipient plasmolysis reveals decreasing turgor pressure in trichoblasts with increasing media stiffness

Water influx is a driver of growth and pushes against an extensible cell wall to help plant cells grow and expand (Cosgrove, 2016; Yi et al., 2022). As turgor pressure increases due to water influx constrained by the cell wall, an excessive rise in turgor pressure beyond the mechanical strength of the cell wall can lead to cell bursting. Based on observing shorter and fewer root hairs in plants grown in increasing media stiffness, we hypothesized that turgor pressure would also follow a negative trend with increasing mechanical stress. To estimate turgor pressure, we modified an incipient plasmolysis method previously described for stomatal guard cells for use in trichoblasts **(Fig. 3A)** (Chen et al., 2021). Plasmolysis of trichoblasts with increasing osmolyte concentrations followed a non-linear regression curve **(Fig. 3B)**. Trichoblasts in roots grown in 0.5% agar media required a higher sorbitol concentration to plasmolyze compared to those grown in 1% and 2% agar media. We solved the regression curves for each dataset and calculated turgor pressure using a modified form of the universal gas equation (ψ = cRT). A decreasing trend in trichoblast turgor pressure was observed with increasing agar concentration in the growth media. Average turgor pressure in trichoblasts grown in 0.5% agar media was 0.86 ± 0.05 MPa, and in 1% and 2% media was 0.77 ± 0.01 MPa and 0.65 ± 0.06 MPa, respectively, and all these values were significantly different from each other **(Fig. 3C)**. The same trend was also observed in atrichoblasts **(Supp. Fig. S3)**. Decreased turgor pressure in trichoblasts further explains the observed phenotype of shorter root hairs with increasing media stiffness.

**Fig 3.**
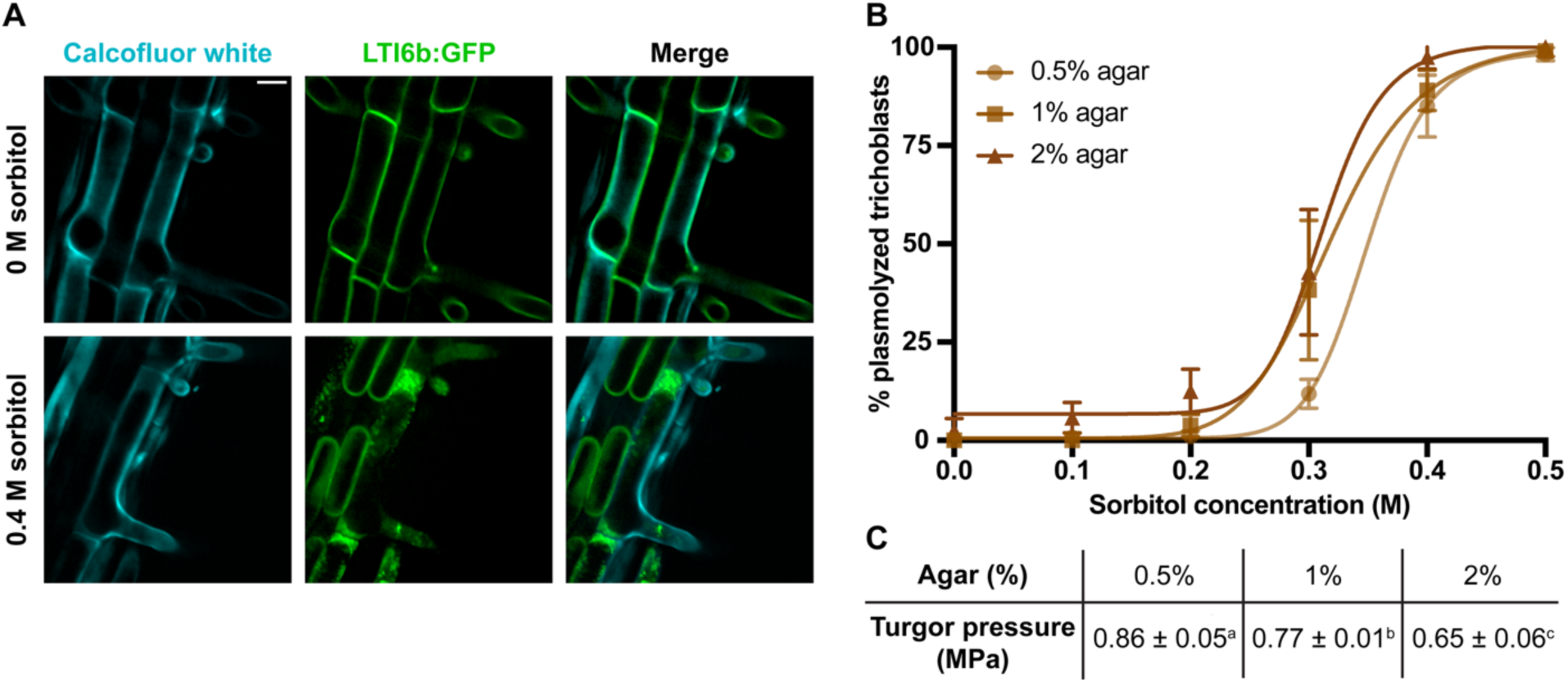
Turgor pressure in trichoblasts decreases with increasing media stiffness. **A.** Representative images of 5-day-old roots expressing LTI6b-GFP grown in media supplemented with 1% agar (Scale bar = 20 µm). Upper row: Seedlings treated with 0 M sorbitol, showing non-plasmolyzed root epidermal cells. Lower row: Seedlings treated with 0.4 M sorbitol, showing plasmolyzed root epidermal cells. Cyan signal represents cell walls stained with Calcofluor white; green signal represents LTI6b:GFP plasma membrane marker. **B,** Non-linear regression curves representing percent plasmolysis of trichoblasts in different sorbitol concentrations. **C,** Turgor pressure in trichoblasts as calculated using the plasmolysis trend. Superscript letters represent statistically significant difference. n = 12-16 seedlings per agar concentration per sorbitol treatment from 5 independent experiments. Different letters denote significance, p < 0.05, ANOVA and post-hoc Tukey test.

### Root hairs grow faster with increasing media stiffness

Since we observed shorter root hairs and decreased turgor pressure with increasing media stiffness **(Fig. 1D and 3C)**, we hypothesized that we would also observe a decrease in the growth rate of root hairs grown into increasing media stiffness.

To allow for high-resolution imaging, we grew seedlings into Ibidi^®^ slides, which are macrofluidic chambers that allow for live cell imaging in a 3D microenvironment **(Fig. 4A)**. Contrary to our initial hypothesis, we observed that root hairs grew faster with increasing agar concentration **(Fig. 4B)**. Mean growth rates in 0.5%, 1%, and 2% agar media were 0.47 ± 0.25 µm/min, 0.62 ± 0.34 µm/min, and 0.81 ± 0.32 µm/min, respectively. To test whether these growth rates were dependent on solid agar media, we repeated these experiments using granular agar and observed a similar trend of increasing growth rate **(Supp. Fig. S4)**. These results contrast with prior observations of decreasing growth rate with increasing media stiffness for root hairs grown in microfluidic chambers (Pereira et al., 2024) and also suggest the existence of changing cell wall mechanics in root hairs under different mechanical environments, since growth rate is determined by turgor pressure, wall yielding, and the rate of deposition of cell wall materials. Therefore, we decided to quantify cell wall properties in root hairs.

**Fig 4.**
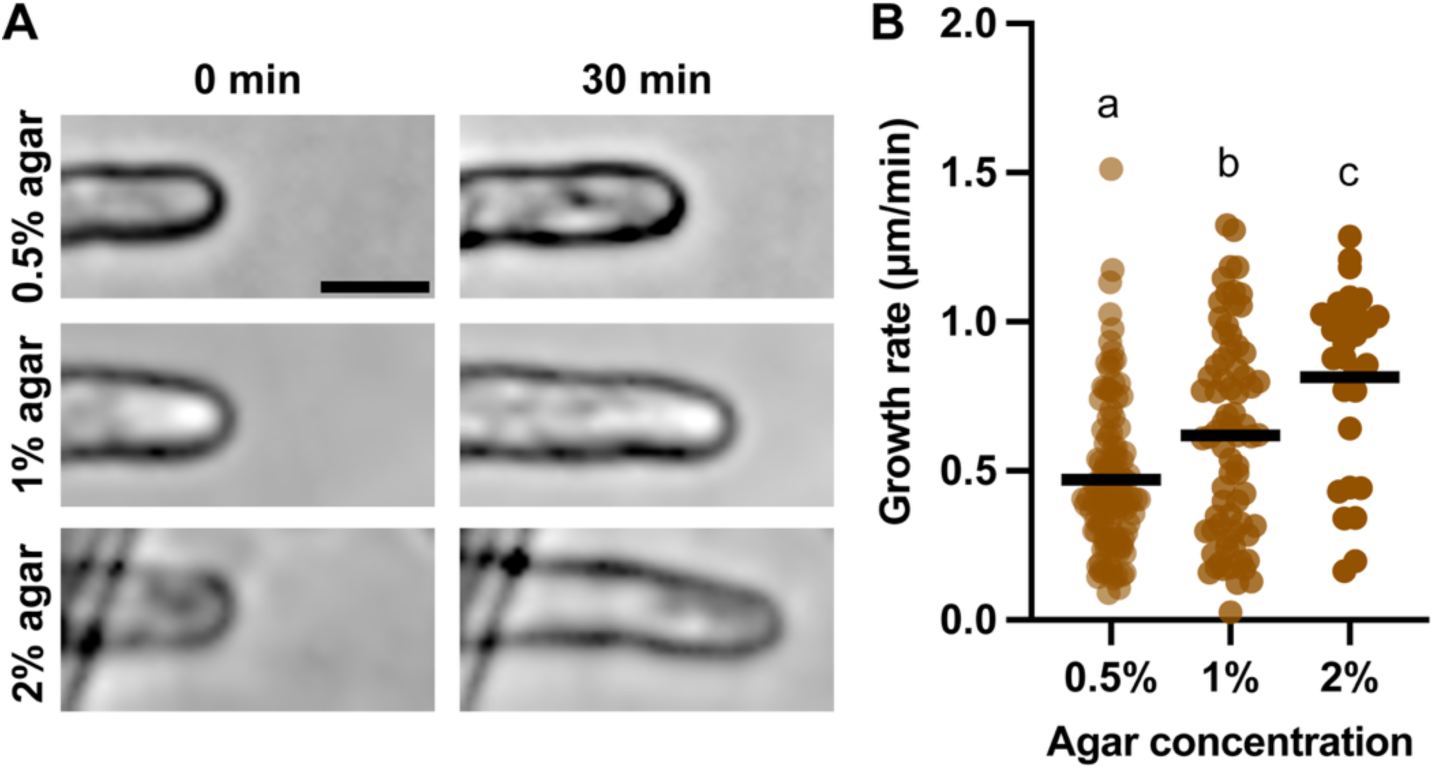
The growth rate of root hairs increases with the stiffness of the growth environment. **A,** Representative brightfield images of root hairs growing in different media at two timepoints, (Scalebar = 20 µm). **B,** Growth rate quantified for 30 min. Each data point indicates one root hair. n = 28-113 root hairs from at least 12 seedlings per growth condition from 5 independent experiments. Different letters denote significance, p < 0.05, ANOVA and post-hoc Tukey test.

### Root hairs have thickened cell walls and cellulose fibrils oriented at larger angles relative to the growth axis of root hairs in the stiffest media

Because cellulose is a major load-bearing component of the cell wall (Paredez et al., 2006), we stained 5-day-old Col-0 seedlings that were grown in mechanically differing media for cellulose using Pontamine Fast Scarlet 4B (S4B) (Anderson et al., 2010) and collected image stacks of mature root hairs **(Fig. 5A and 5C)**. Quantification of cell wall thickness at multiple locations along root hair shanks revealed thicker cell walls in the stiffest growth medium **(Fig. 5A)**. Wall thicknesses of root hairs grown in 0.5%, 1%, and 2% agar media were 448.5 ± 48.55 nm, 472.4 ± 58.35 nm, and 567.6 ± 103.9 nm, respectively **(Fig. 5B)**. Simultaneously, we quantified root hair diameter and found an increased diameter in root hairs grown in 1% and 2% agar, with 11.71 ± 1.34 µm and 11.46 ± 3.37 µm diameters respectively, as compared to root hairs grown in 0.5% agar with a 10.45 ± 1.27 µm diameter **(Fig. 5C)**. The cross-sectional areas and perimeters of root hairs followed mixed trends **(Supp. Fig. S5)**. Root hair area in 0.5% agar was 98.02 ± 24.88 µm^2^ but was significantly larger in root hairs grown in 1% agar at 125.6 ± 32.08 µm^2^. Root hair area in 2% agar was 110.7 ± 51.89 µm^2^, which was not significantly different from root hairs grown in 0.5% or 1% agar media **(Supp. Fig. S5A)**. Root hair perimeter also followed a mixed trend. The average perimeters of root hairs grown in 0.5% and 2% agar media were 34.91 ± 4.27 µm and 36.38 ± 8.49 µm, respectively, and did not differ significantly, but root hair perimeter was significantly higher in 1% agar at 39.56 ± 4.96 µm **(Supp. Fig. S5B)**. We also quantified cellulose microfibril orientation and anisotropy in root hairs **(Fig. 5D)**. Cellulose angle was larger with respect to the growth axis in root hairs grown in 2% agar at 32.92 ± 21.74 degrees than in root hairs grown in 0.5% and 1% agar, where cellulose orientation was 20.84 ± 17 and 19.66 ± 17.53 degrees, respectively **(Fig. 5E)**. The optical anisotropy in S4B staining in root hairs remained unchanged across different treatments **(Fig. 5F)**. We speculate that the larger cellulose angle in the stiffest media might reflect a response mounted by root hairs to reinforce the circumferential wall of the root hair, favoring cell elongation against higher mechanical impedance but potentially resulting in earlier growth cessation.

**Fig 5.**
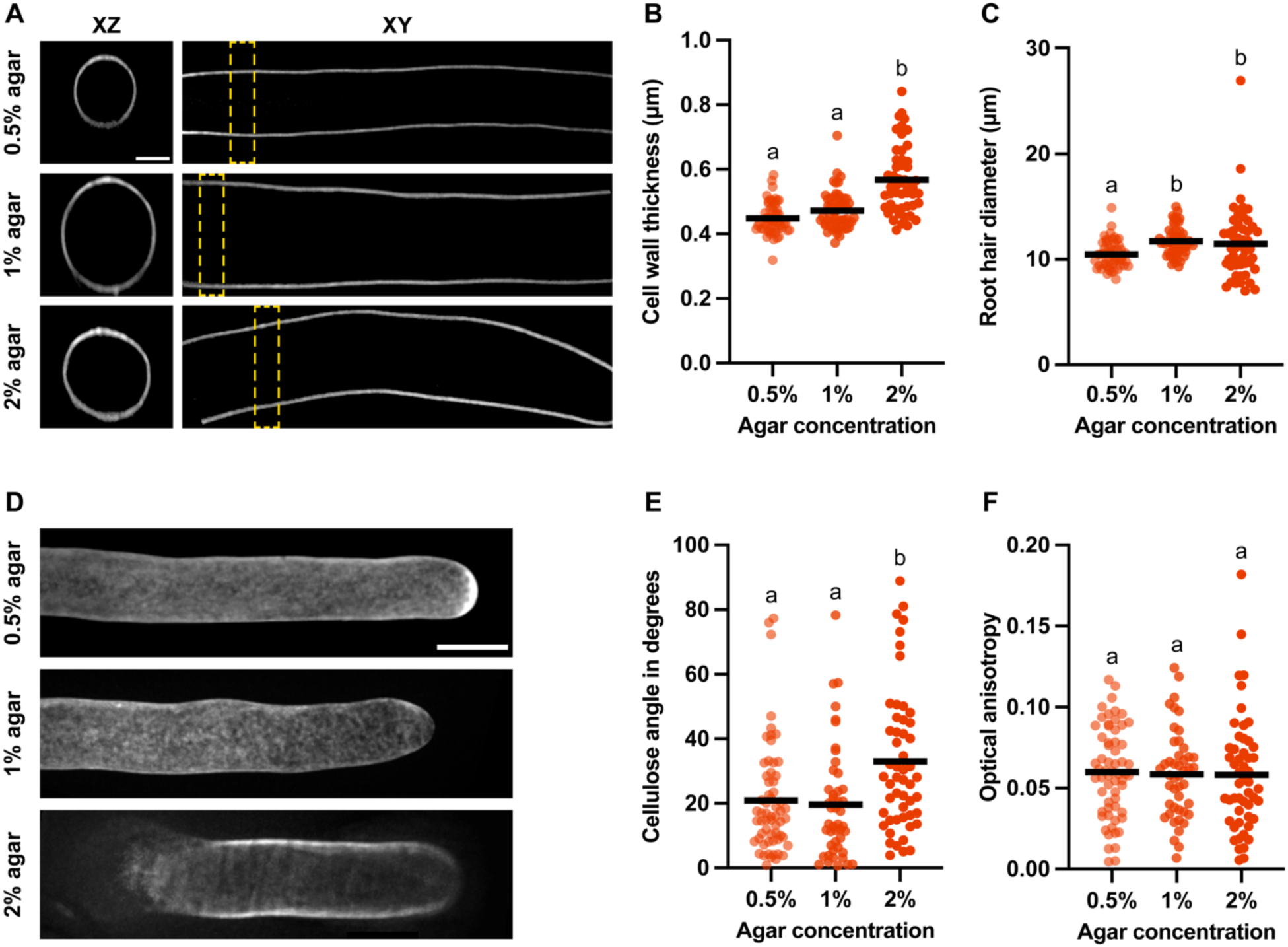
Cell wall thickness, root hair diameter, and cellulose orientation show highest magnitudes in the stiffest growth environment. **A,** 5-day-old Col-0 root hairs grown in mechanically distinct media and stained with S4B for cellulose (Scale bar = 5 µm). Images show XZ and XY cross-sections of root hairs. Dashed yellow boxes on XY sections show regions of corresponding XZ projections. **B,** Cell wall thickness as measured by the FWHM method. **C,** Root hair diameter as measured by plotting a fluorescent intensity profile across the root hair and measuring the distance between the intensity peaks on either side of the root hair. Diameters were measured at three locations on a root hair shank, and the average was plotted per root hair. **D,** Periclinal surfaces of 5-day-old Col-0 root hairs stained for cellulose (Scale bar = 10 µm). **E,** Net cellulose orientation in individual root hair shanks as quantified by FibrilTool. **F,** Optical anisotropy as quantified by FibrilTool. Each data point indicates one root hair. n ≥ 51 root hairs from at least 15 seedlings per growth condition from 5 independent experiment. Different letters denote significance, p < 0.05, ANOVA and post-hoc Tukey test.

### Microtubule orientation follows a similar pattern as microfibrils, but at different angles, and microtubule density and anisotropy decrease in root hairs with increasing media stiffness

Microtubules are crucial for regulating the directionality and stability of root hair growth, as their depolymerization causes aberrant root hair phenotypes (Bibikova et al., 1999). Additionally, microtubules guide cellulose synthase complexes (CSCs) to pattern cell walls (Paredez et al., 2006) and are sensitive to changes in the mechanoenvironment (Xiao et al., 2016). Therefore, we quantified microtubule orientation in root hairs grown in different agar percentages. Microtubules labeled with GFP-MAP4 followed a similar trend in orientation as cellulose, but at smaller angles with respect to the growth axis of root hairs **(Fig. 6A)**. Average microtubule angle in root hairs grown in 0.5%, 1%, and 2% agar media was 4.87 ± 3.85, 4.5 ± 3.56, and 7.42 ± 6.12 degrees, respectively **(Fig. 6B)**. These orientation values were similarly proportionately lower in 0.5%, 1%, and 2% agar media than microfibril orientations, which were 4.28, 4.37, and 4.44 times the microtubule angles, respectively **(Fig. 5E)**. We also observed a decremental trend in microtubule anisotropy **(Fig. 6C)**. Microtubule anisotropy was 0.34 ± 0.12 in root hairs grown in 0.5% agar, 0.31 ± 0.11 in root hairs grown in 1% agar, and 0.28 ± 0.15 in root hairs grown in 2% agar, which was significantly lower than 0.5% agar treatment. This disruption in microtubule organization might impair anisotropic cell expansion, resulting in shorter root hairs. Microtubule density, as measured by the area of microtubules occupied per unit area of root hair, also decreased with increasing agar concentration **(Fig. 6D)**. Decreased microtubule density might make it harder to maintain root hair polarity and hence expansion.

**Fig. 6.**
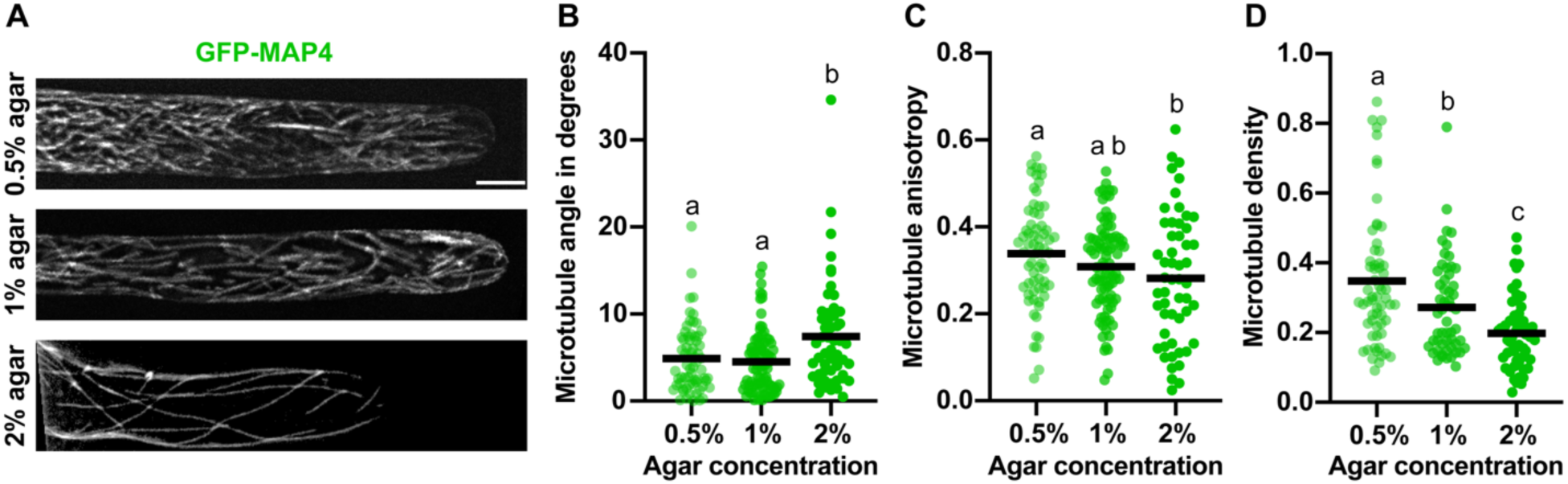
Microtubules become more transverse relative to the growth axis, whereas their anisotropy and density decrease, in root hairs grown under increasing mechanical stress. **A,** Representative images of 5-day-old root hairs expressing GFP-MAP4 grown in increasing concentrations of agar. Fluorescence represents microtubules. (Scale bar = 10 µm). **B,** Net microtubule orientation relative to the growth axis as quantified by FibrilTool. **C,** Image anisotropy as quantified by FibrilTool. **D,** Microtubule density as represented by the percentage of microtubule area with respect to root hair area. Each data point represents a single root hair. n ≥ 53 root hairs from at least 18 seedlings per growth condition from 5 independent experiments. Different letters denote significance, p < 0.05, ANOVA and post-hoc Tukey test.

### Finite element models of root hair growth predict lower surface stress with simulated increases in media stiffness

Finite Element Modeling (FEM) can predict the behaviors of a specific geometry under varying mechanical conditions, including pressure, external and internal constraints, and material properties. FEM has been successfully used to study the mechanical properties of biological systems like cells and tissues, yielding detailed insights into their growth patterns and interactions with their microenvironments (Bidhendi & Geitmann, 2018; Yi et al., 2022). Differences in the mechanical stiffness of the growth environment affect turgor pressure and cell wall properties in root hairs and alter their growth **(Figs. 3-5)**. These phenotypes could potentially be mediated by changes in wall stress, which can activate mechanical signaling pathways (Duan et al., 2010; Wolf et al., 2012).

To predict surface stress in root hairs, we simulated FEMs with average wall thicknesses and turgor pressures observed under different agar conditions **(Fig. 5B)** and constant longitudinal (E1) and circumferential (E2) moduli estimated with the average cellulose orientations observed in root hair shanks **(Fig. 5E)**. We also ran sensitivity analysis by swapping magnitudes of one variable at a time and keeping the other two variables constant to determine which factor amongst turgor pressure, wall thickness, and wall modulus had the highest impact on the surface stress. Sensitivity analysis was followed by Relative Weights Analysis (RWA) using the rwa() R package. RWA showed that the surface stress on root hair FEMs is mainly explained by variation in turgor pressure **(Supp. Table S1)**. Therefore, we ran root hair FEMs with individual values of turgor pressures observed in independent experiments **(Supp. Table S2)** paired with average cell wall thicknesses and circumferential moduli of root hairs from the different agar treatments. The initial state of FEM was at 0 MPa surface stress **(Fig. 7A)**. After simulating growth, we found that surface stress on root hair FEMs was inversely proportional to agar concentration **(Fig. 7B-D)**. The highest surface stress in the root hair FEM with 0.5% agar parameters was 11.85 ± 0.56 MPa, whereas with 1% agar parameters it was 10.23 ± 0.14 MPa, and in 2% agar parameters it was 7.5 ± 0.63 MPa **(Fig. 7E)**. We also observed that surface stresses at root hair tips were always lower than their corresponding shank regions. The lowest surface stress of 3.18 MPa was seen at the root hair tip in the FEM run under 2% agar parameters **(Fig. 7D)**. This further suggests the ability of the cell wall to yield under limited turgor pressure and promote rapid cell expansion.

**Fig. 7.**
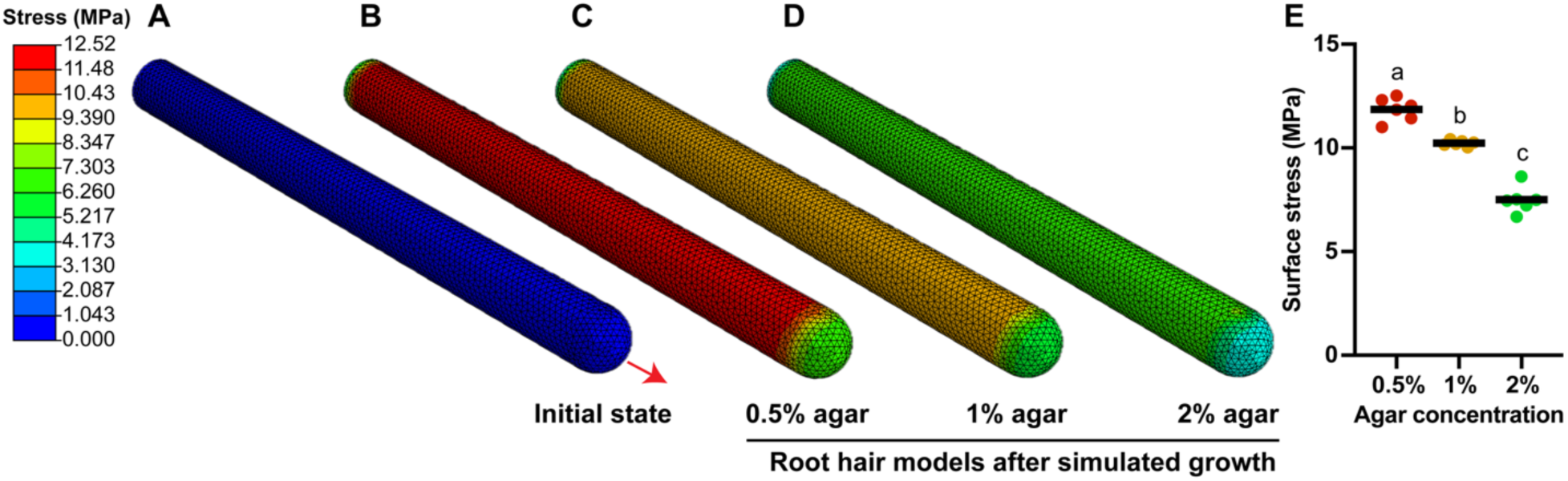
Finite Element Modeling predicts a reduction in surface stress for elongating root hairs grown in media with increasing stiffness. **A,** Initial state of root hair FEM before simulation. Red arrow indicates growth direction. **B-D,** Representative *in silico* finite element models of root hairs run from an initial state defined using corresponding experimental values of cell wall thickness (Fig. 5) and turgor pressure (Fig. 3) for different media stiffnesses. The simulated elongation represents the growth for 10 minutes of the experimental observation. **E,** Maximum surface stress in individual *in silico* root hair models; n = 6 models per condition run with turgor pressure measured in independent *in vivo* experiments. Different letters denote significance, p < 0.05, ANOVA and post-hoc Tukey test.

## Discussion

Many studies have done phenotypic characterization and have quantified the subcellular and extracellular factors that influence root hair growth, such as microtubule and cell wall dynamics, turgor pressure, and Young’s modulus, individually, in the absence of mechanical constraints (Herburger et al., 2022; Lew, 1996; Park et al., 2011; Pereira et al., 2023, 2024; Van Bruaene et al., 2004). Much less information is available regarding the mechanobiological effects of the growth environments on root hair growth and their subsequent cellular and subcellular responses. Here, we identified phenotypic, cellular, and mechanical alterations in root hairs experiencing increasing mechanical perturbation, which resembles their natural growth in soils of varied stiffness.

We started by growing Col-0 seedlings into growth media with increasing agar stiffness and observed that higher media stiffness resulted in shorter root hairs (**Fig. 1D**). A similar trend was observed when root hairs were grown in increasing concentrations of agar in microfluidic devices (Pereira et al., 2024). Pereira et al. used agar concentrations of 0.5%, 1%, and 1.25% instead of 0.5%, 1%, and 2% in this study. Their measurements of average root hair length were 498 ± 20 µm and 339 ± 12 µm in 0.5% and 1% agar, respectively, which were longer than observed in this study **(Fig. 1D).** Additional differences, such as sucrose concentration and growth conditions, between our studies might have led to differences in root hair lengths. Indeed, root hairs exhibit substantial phenotypic plasticity (Müller & Schmidt, 2004; Salazar-Henao et al., 2016). Nonetheless, the trend in decreasing root hair length with increasing media stiffness was similar between the studies. We also observed lower root hair density in response to increasing mechanical stiffness in plants **(Fig. 1G)**. Besides epidermal cell patterning **(Fig. 2)** and turgor pressure **(Fig. 3)**, which influence root hair density, several other factors, such as cell wall loosening by xyloglucan endotransglucosylase/hydrolases (XTHs), Expansins 7 and 18, actin and microtubule reorganization at the root hair bulge site, and local cell wall acidification could be affected by changes in media composition (Baluka et al., 2000; Bibikova et al., 1998; Cho & Cosgrove, 2002; Emons & Derksen, 1986; Lin et al., 2011; Vissenberg et al., 2001). It will be interesting to examine how these factors change under mechanical perturbation in future studies.

Pressure probe measurements indicate that turgor pressure in 7-14-day-old Arabidopsis root hairs is 0.68 ± 0.2 MPa (Lew, 1996). In those experiments, seedlings were grown on 0.9 – 1% gellan gum-based media. Nonetheless, this pressure range is similar to the turgor pressure values quantified in trichoblasts using incipient plasmolysis here **(Fig. 3)**. We found that turgor pressure, which is an isotropic stress (Ali et al., 2023), decreased in trichoblasts with increasing mechanical stiffness of the growth media. Initially, we predicted increased turgor in root hairs grown in stiffer media in order to allow them to grow against increased mechanical resistance, but this increased turgor might override the mechanical integrity of the root hair and lead to bursting, similar to what has been observed in lily pollen tubes after artificially increasing turgor pressure (Benkert et al., 1997) and in the root hairs of wall integrity-sensing and wall biosynthesis mutants (Brost et al., 2019; Duan et al., 2010; Favery et al., 2001; Tan et al., 2020). In our experiments, turgor pressure in atrichoblasts also decreased with increasing agar stiffness and was always lower than the corresponding turgor pressure in trichoblasts **(Supp. Fig. S3)**. Still, atrichoblasts were longer than trichoblasts **(Supp. Fig. S1)**, (Beemster & Baskin, 1998; Löfke et al., 2013), which further explains the complex nature of turgor mediated cell expansion. Taken together, our turgor pressure results further explain shorter and fewer root hair phenotypes in stiffer media.

A recent publication also quantified growth rates of root hairs in media with increasing stiffness and observed a decreasing trend (Pereira et al., 2024). The authors also concluded that the growth duration of the root hairs remained the same irrespective of the growth media. In contrast to these results, we observed faster growth in root hairs grown under increasing media stiffness **(Fig. 4)** and **(Supp. Fig. S4)**, which, in combination with our finding that root hairs were shorter with increasing media stiffness **(Fig. 1D)**, implies that the duration of root hair growth decreases. Additionally, rapid wall deposition and modification at root hair tips might lead to faster growth under limited turgor pressure (Galway et al., 1999; Grierson et al., 2014; Monshausen et al., 2007). Actively monitoring cell wall dynamics in root hairs is possible thanks to metabolic click labeling (Anderson et al., 2012; Herburger et al., 2022; McClosky et al., 2016; Mravec et al., 2017), but is technically challenging to perform when root hairs are growing inside agar.

Despite these challenges, we could still measure cellulose patterning and cell wall thickness in root hairs grown in different media **(Fig. 5)**. Cellulose deposition in root hairs is mediated by CELLULOSE SYNTHASE-LIKE D proteins, COBRA-LIKE proteins, and CELLULOSE SYNTHASE proteins (Bernal et al., 2008; Chen et al., 2016; Li et al., 2022; Park et al., 2011; Singh et al., 2008). Though we know about the proteins involved in cellulose synthesis and deposition in the cell walls of root hairs, as mentioned earlier, less information is available about cellulose patterning and cell wall thickness, and how these factors influence root hair expansion (Akkerman et al., 2012; Mendrinna & Persson, 2015). Study of wall texture found that Arabidopsis root hairs have disorganized cell walls at their tips and axially/helically organized walls in their shanks (Akkerman et al., 2012). Cellulose orientation influences cell expansion, and dynamic cellulose reorientation was observed in elongating root epidermal cells (Anderson et al., 2010). Based on our observations of root hair lengths in different agars **(Fig. 1D)**, we expected that cellulose orientation in stiffer media would be more aligned with the growth axis, restricting cell elongation, but on the contrary, we observed steeper angles of cellulose fibrils with respect to the growth axis of root hairs grown in the stiffest media **(Fig. 5D and 5E)**. We speculate that this change in cellulose orientation could be a response mounted by root hairs against mechanical stress to promote axial expansion in the stiffest medium (Chan et al., 2010), and further explains the increased growth rate we observed in root hairs grown in stiffer media.

We found that cell wall thickness, as measured in different regions of root hair shanks, increased with mechanical stress **(Fig. 5A and 5B)**. In a cellulose mutant of *Lotus japonicus*, *Ljclsd1-1*, shorter root hairs with thicker cell walls were observed (Majda, 2021). In light of the anisotropic nature of the cell wall, thicker walls in root hairs might restrict radial expansion and promote axial expansion, further explaining the increased growth rate we observed in root hairs **(Fig. 4)** and **(Supp. Fig. S4)**. Although root hair diameter slightly increased with increasing media stiffness **(Fig. 5C)**, cross-sectional perimeter and area in root hairs did not differ significantly across different media stiffnesses **(Supp. Fig. S5)**. Altogether, the changes we observed in cell wall thickness and cellulose orientation under increasing media stiffness imply that root hairs modulate wall patterning in response to the mechano-environment to maintain the capacity for growth, but that physical limitations imposed by the growth environment, in combination with mechanically induced signaling, might place limits on the ability of the cell to adapt by changing cell wall patterning.

Microtubules are essential for establishing and maintaining cell polarity in root hairs and directing cell growth. They accomplish this by forming a framework that guides vesicle delivery, streamlines nuclear movement, and regulates the orientation of cellulose deposition (Brueggeman et al., 2022; Chan et al., 2010; Sieberer et al., 2005). All these processes are essential for normal growth in root hairs. One study in cotyledon pavement cells found that microtubules reorient parallel to the direction of mechanical stress (Sampathkumar et al., 2014). This finding demonstrates the mechanosensitive behavior of microtubules. In our study, root hairs experienced uniform stress throughout their growth period from all directions, which led to a net perpendicular orientation of microtubules with respect to the axis of root hair growth in the stiffest media **(Fig. 6A and 6B)**. A major reason for the disparity between the magnitudes of cellulose microfibril and microtubule angles might be due to the activity of cellulose synthesis proteins like CELLULOSE SYNTHASE-LIKE D isoforms being independent of microtubule tracks, as previously reported in *Arabidopsis thaliana* and *Physcomitrium patens* (Wu et al., 2023; Yang et al., 2020). Additionally, decreased microtubule anisotropy **(Fig. 6C)** and density **(Fig. 6D)** imply that the localized delivery of cell wall material might be disrupted under high mechanical stiffness in the growth environment, resulting in both aberrant wall patterning and a shorter duration of root hair growth.

Finally, we built and tested FEMs of root hair to study the cell wall stress in mechanically different growth environments **(Fig. 7)**. Analogous FEMs of pollen tubes were previously built to study polar cell growth, (Bidhendi & Geitmann, 2018; Fayant et al., 2010), but root hairs display important distinctions in terms of wall structure and growth dynamics from pollen tubes (Herburger et al., 2022; Mendrinna & Persson, 2015). An anisotropic-viscoplastic model was proposed to explain the anisotropic surface expansion in the root hairs of *Medicago truncatula* (Dumais et al., 2004, 2006). However, this mechanical model only considered the tip growth of root hairs. Latest updates show wall expansion in the shank region of root hairs as well (Herburger et al., 2022). Our FEMs favored anisotropic expansion of root hairs, which was also suggested to be the best fit to explain polar growth in root hair tips (Dumais et al., 2004). Our root hair FEM predicted higher stress in the shank region compared to the tips of root hairs under all growth conditions **(Fig. 7)**. This resembles the stress profiles previously reported in FEMs of lily pollen tubes (Fayant et al., 2010) suggesting a mechanically stiffer cell wall in the shank than in the tip region. Our FEMs also predict an overall reduction in the cell wall stress of root hairs with increasing media stiffness and further explain the observed growth phenotypes observed under different mechanical constraints **(Figs. 1, 3, and 4)**. A future challenge is to develop a realistic model of root hair growth incorporating anisotropic polar growth, mediated by the simultaneous actions of turgor pressure and wall loosening, and also to simulate wall synthesis and deposition at both the tip and the shank.

In summary, our data depict the morphogenetic, subcellular, and biomechanical changes in root hair growth in *Arabidopsis thaliana* in response to mechanical cues. Based on this foundation, further studies can explore the specific genetic and molecular components involved in cell wall integrity sensing and mechanotransduction pathways in root hairs. This will help us understand how root hairs optimize their growth under challenging soil conditions and, in the long run, engineer plants that can sustain soil hardening.

## Materials and methods

### Plant materials and growth conditions

*Arabidopsis thaliana* (Col-0) was used for studying root hair morphogenetics, epidermal cell patterning, analyzing the growth rates of root hairs, and quantifying cellulose orientation. For incipient plasmolysis experiments, a membrane marker line (LTI6b:GFP) was used (ABRC Stock No. CS84726), and for imaging microtubule dynamics, GFP-MAP4/Col-0 line was used (Marc et al., 1998). Seeds were surface sterilized with 30% bleach and 0.1% SDS and washed with sterile Mili-Q^®^ four times and suspended in 0.15% agar solution. Subsequently, they were stratified in the dark at 4°C for at least 3-days. All the Arabidopsis lines were grown in ½ MS media (Calsson labs Murashige and Skoog basal salts MSP01), 0.06% (w/v) MES, pH adjusted to 5.6, and supplemented with 1% sucrose and 0.5%, 1%, and 2% agar (Sigma Aldrich A4550, CAS-No:9002-18-0) in 2 ml microcentrifuge tubes (Globe Scientific Inc.) for 5-days before imaging. Seeds were gently sown slightly beneath the top surface of the media, and 2-3 holes were poked into the caps of microcentrifuge tubes using sterile needles (18G x 1 ½”, Air-Tite Products Co. Inc.). Punched tubes were covered with Micropore^TM^ tape (3M Co.) to facilitate air exchange and avoid contamination. Tube racks were placed in the plant growth chamber (Percival) at 24-hour light setting with light intensity of 120-150 μmol/m²/s.

### Root hair growth rate measurements

Col-0 seedlings were grown in Ibidi^®^ µ-Slide VI 0.4 Uncoated (Cat.No: 80601) filled with sterile ½ MS growth media supplemented with 1% sucrose and 0.5%, 1%, and 2% agar respectively. For corroboration experiments with granular media, high gelling temperature agarose (ThermoScientific Cat.No: J66704.22, CAS#: 9012-36-6) was dissolved in deionized water at 85°C and autoclaved at 121°C for 20 mins. It was cooled while stirring on a magnetic stirrer at 900 RPM overnight. These granular agars were mixed with liquid growth media such that the final concentration of media was ½ MS with 1% sucrose. These agar layers were injected into Ibidi^®^ µ-Slide channels to grow seedlings for live cell imaging. Growing root hairs were imaged using bright field Z-stack combined with time lapse every minute for a duration of 30 minutes. To quantify the growth rates of root hairs, movies were analyzed in Fiji (Schindelin et al., 2012).

### Incipient plasmolysis to calculate turgor pressure

5-day-old LTI6b-GFP seedlings were gently pulled out of the growth media in the individual microcentrifuge tubes and transferred to sorbitol solutions of differing concentrations (0, 0.1, 0.2, 0.3, 0.4, and 0.5 M) for at least 6 mins and subsequently mounted on microscope slides (VWR^®^ VistaVision^TM^ Cat.No. 16004-430) with double-sided tape chamber containing sorbitol solution with respective concentration (∼ 2 mins). The slide was further mounted to the microscope stage (Zeiss Axio Imager.M2) and root hairs and epidermal cells (trichoblasts and atrichoblasts) were focused using 40X oil immersion objective (∼ 2 mins). The number of plasmolyzed and non-plasmolyzed cells were counted for the next 5 mins and reported as the percentage plasmolysis and plotted as a function of increasing sorbitol concentration. A non-linear regression curve was fitted, and equations were solved to find the sorbitol concentration required to plasmolyze 50% of the epidermal cells observed. Corresponding sorbitol concentrations were substituted in the modified form of the universal gas equation ψ = cRT, where ψ is the turgor pressure in Pascals, c is the sorbitol concentration required to plasmolyze 50% of the epidermal cells, R is the universal gas constant (8.3145 J⋅K⁻¹⋅mol⁻¹), and T is the average room temperature of the day in Kelvin.

### Confocal microscopy

We used Zeiss spinning disk confocal microscope (Observer.Z1) with a Yokogawa CSU-X1 spinning head. Seedling was mounted on 3×1×1 glass slides (VWR^®^ VistaVision^TM^ Cat.No. 16004-430) within a small chamber that was made using double sided stick-tape (3M^TM^ Scotch^®^ Double Sided Tape 665), and in 50-60 µl mounting solution as mentioned below. A 24×40 mm rectangular #1½ cover glass (Corning^®^ Cat.No. 2980-244) was placed on the seedling but resting on the tape making sure not to squish the seedling. Zeiss immersion oil (Immersol^TM^ 518 F) was applied to the cover glass when using oil-immersion objectives. Confocal images were deconvoluted using AutoQuantX2 software.

#### Propidium iodide staining

5-day-old Col-0 seedlings were incubated in dark for 10 minutes in 10 µg/ml propidium iodide solution (Sigma Aldrich P4864) prepared in Milil-Q^®^. Seedlings were washed twice with Mili-Q^®^, mounted in Mili-Q^®^ on glass slide, and imaged using 20X objective by taking Z-stack series with step distance of 1 µm. 561 nm excitation laser with 617/73 nm emission filter was used to detect propidium iodide fluorescence. These confocal images were used to quantify epidermal cell patterning and their lengths.

#### Calcofluor white staining

5-day-old LTI6b:GFP seedlings were incubated in dark for 10 minutes in 1% calcofluor white solution prepared in Mili-Q^®^. Seedlings were washed once with Mili-Q^®^, mounted in Mili-Q^®^ on glass slide, and imaged under 40X oil immersion objective by taking Z-stack series with step distance of 0.5 µm. Calcofluor white binds to β-1,3- and β-1,4-linked glucans and fluoresces when excited with 405 nm laser, while GFP tagged to LTI6b, a membrane protein, fluoresces when excited with 488 nm laser and fluorescence was detected using 450/50 nm and 525/50 nm emission filters respectively. Merged fluorescent images helped distinguish plasmolyzed epidermal cells from non-plasmolyzed cells to calculate turgor pressure.

#### Pontamine fast scarlet 4B (S4B) staining

5-day-old Col-0 seedlings were incubated in dark for 30 minutes in 0.01% S4B solution prepared in liquid ½ MS medium as described previously (Anderson et al., 2010). Seedlings were washed once in ½ MS liquid medium, mounted in ½ MS on glass slide, and imaged under 100X oil immersion objective by taking Z-stack series with step distance of 0.2 µm. S4B stains cellulose and fluoresces when excited with 561 nm laser and fluorescence was detected using 617/73 nm emission filter. These confocal images were used to quantify cellulose orientation and cell wall thickness in root hairs.

#### Microtubule imaging

5-day-old GFP-MAP4 seedlings were gently pulled out of their respective microcentrifuge tubes and directly mounted on slide in Mili-Q^®^. Root hairs were imaged under 63X oil immersion objective by taking Z-stack series with step distance of 0.2 µm. GFP tagged to MAP4, a microtubule associated protein, fluoresces when excited with 488 nm laser and the fluorescence was detected using 525/50 nm emission filter. These confocal images were used to quantify microtubule orientation, density, and anisotropy in mature root hairs.

### Image analysis and quantification

Confocal image analysis was performed with Fiji using inbuilt tools and plugins (Schindelin et al., 2012). Root hair morphogenetics analysis, such as length and number quantification, was done using the segmented line tool and the multipoint tool, respectively. Line tool was also used to quantify root hair growth rates from the time lapse movies. Epidermal cell patterning was analyzed using BigTrace 0.4.1 plug-in that helps render 3D image for faster cell tracing. Cellulose microfibril and microtubule orientations and anisotropy were quantified using an ImageJ plug-in FibrilTool (Boudaoud et al., 2014). Cell wall thickness was quantified by plotting the fluorescent intensity profile followed by the Full Width Half Max (FWHM) method (Komis et al., 2015). Microtubule density was calculated by quantifying the total area occupied by microtubules inside root hairs using image thresholding and dividing it by the total area of root hair in the image frame.

### Finite Element Modeling

For the computational simulation of the root hair growth using the Finite Element method, the root hair was modeled as a hollow cylinder with a 5 μm radius capped with a half-sphere. This geometric model was meshed using Gmsh (v 4.15) and imported into Abaqus (v. 2023). The imported root hair mesh was modeled as a shell with an appropriate thickness value reflecting experimental observation. The growth of root hair was modeled as a static mechanical response to the uniform turgor pressure applied to the inner surface, which corresponds to the growth of root hair observed in experiments over 10 min. A linear anisotropic elasticity plane-strain was assigned uniformly across the whole cell wall. Reflecting the orientations of cellulose, E1 was defined as longitudinal modulus, and E2 was defined as circumferential modulus, respectively.

### Statistical analysis

Statistical analyses were performed and graphs were plotted with GraphPad v10.4.0. Individual data points are represented by dots, while horizontal lines indicate the mean value. Details regarding sample size, statistical significance annotations, and the statistical tests used for each plot are provided in the figure legends.

## Acknowledgements

Thanks to Gavin McDermond for help with measurements, and Dr. Megan Cooke and Dr. Jason Burdick (University of Colorado Boulder) for providing granular agar media.

## Author contributions

VZ and CTA designed experiments, VZ performed experiments and analyzed data, VZ and CTA interpreted data, HY and VZ built and tested the FEM, VZ drafted the manuscript, and all authors reviewed and edited the manuscript.

## Funding

This work is supported by the NSF Center for Engineering Mechanobiology (CEMB), NSF grant CMMI-154857, and a CEMB Trainee Pilot Award to Vyankatesh Zambare.

## Notes

### Competing Interest Statement

The authors have declared no competing interest.

