## Supplementary Material for "Mechanical constraint causes lower turgor, thicker walls, and faster growth in Arabidopsis root hairs"

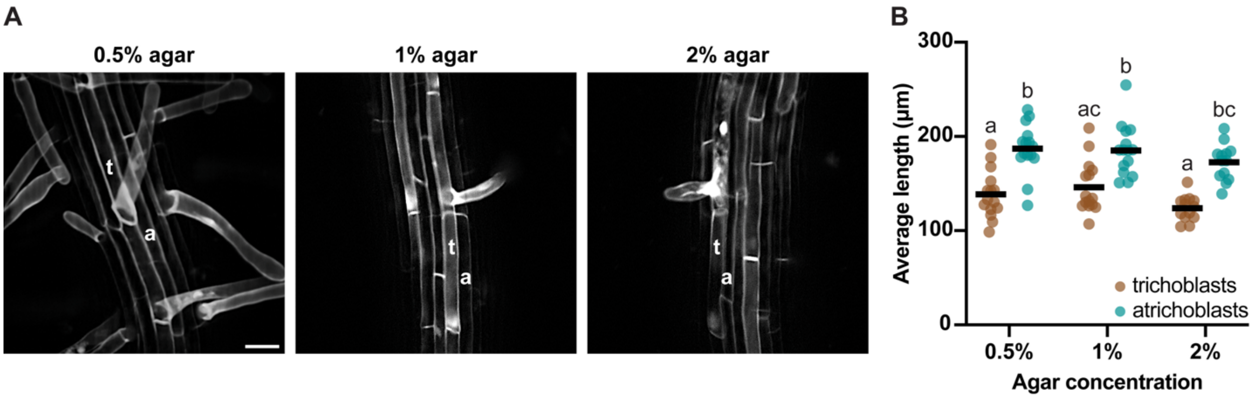


**Supplementary Fig. S1. Increasing media stiffness does not affect the lengths of root epidermal cells**

**A,** Maximum projections of 5-day-old Col-0 roots grown in media with different agar concentrations and stained with propidium iodide. t-trichoblast, a-atrichoblast (Scale bar = 40 µm). **B,** Lengths of trichoblasts and atrichoblasts do not vary significantly in different agar concentrations, and atrichoblasts are longer than trichoblasts in all conditions. $n \geq 13$ seedlings from 5 independent experiments. Different letters denote significance, $p < 0.05$, ANOVA and post-hoc Tukey test.

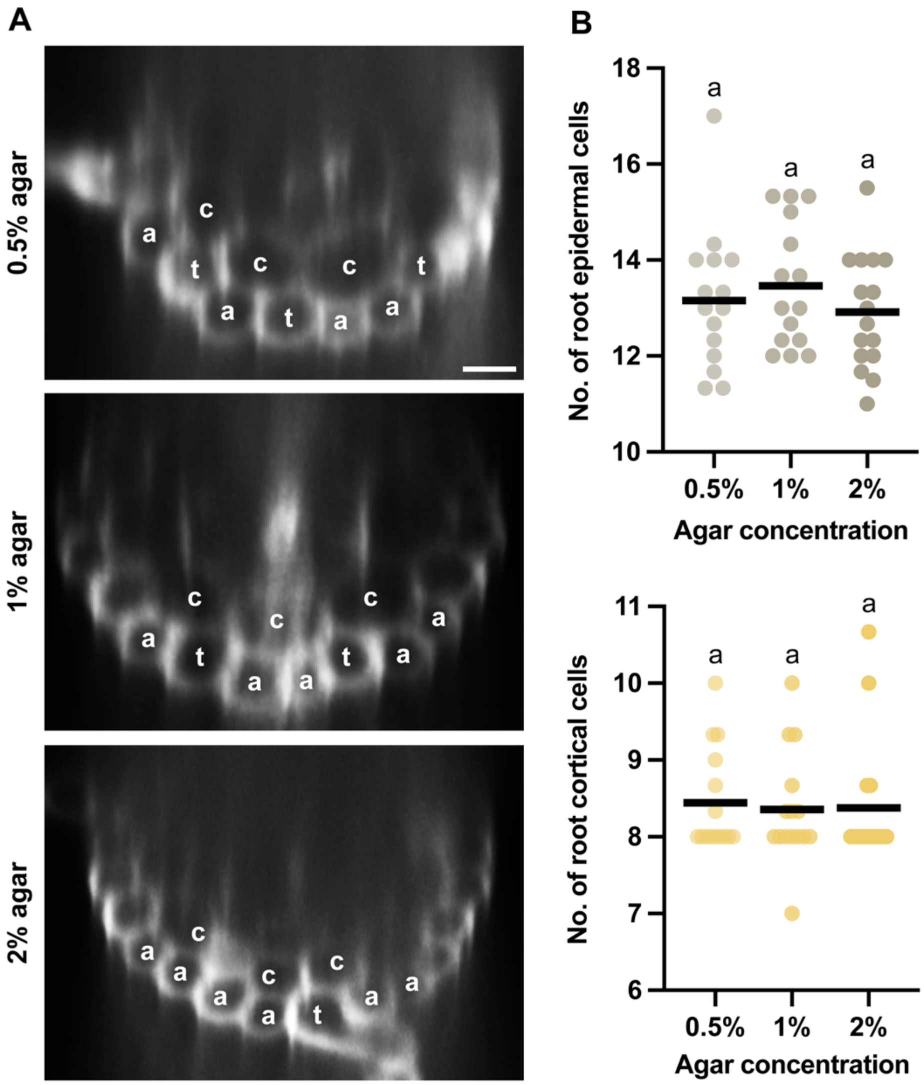


**Supplementary Fig. S2. Increasing media stiffness perturbs root epidermal cell patterning without changing the total number of epidermal and cortical cells**

**A,** XZ cross-sections of 5-day-old Col-0 roots grown in media with different agar concentrations and stained with propidium iodide. c-cortical cell, t-trichoblast, a-atrichoblast (Scale bar = 20 µm). **B,** Numbers of root epidermal and cortical cells do not vary significantly in different agar concentrations. $n \geq 15$ seedlings from 6 independent experiments. Different letters denote significance, $p < 0.05$, ANOVA and post-hoc Tukey test.

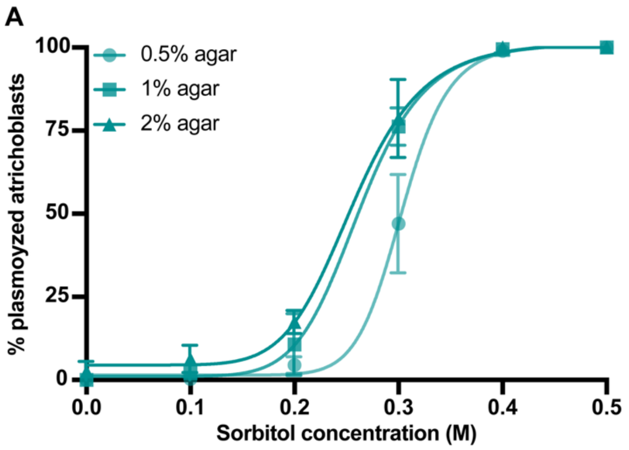


B

| Agar (%) | 0.5% | 1% | 2% |
|---|---|---|---|
| Turgor pressure (MPa) | $0.71 \pm 0.06^{a}$ | $0.62 \pm 0.05^{b}$ | $0.55 \pm 0.02^{b}$ |

**Supplementary Fig. S3. Turgor pressure in atrichoblasts decreases with increasing media stiffness**
**A,** Non-linear regression curves representing percentage plasmolysis of atrichoblasts in different sorbitol concentrations. **B,** Turgor pressure in atrichoblasts as calculated using the plasmolysis trend. n = 12-16 seedlings per agar concentration per sorbitol treatment from 5 independent experiments. Different superscript letters denote significance, $p < 0.05$, ANOVA and post-hoc Tukey test.

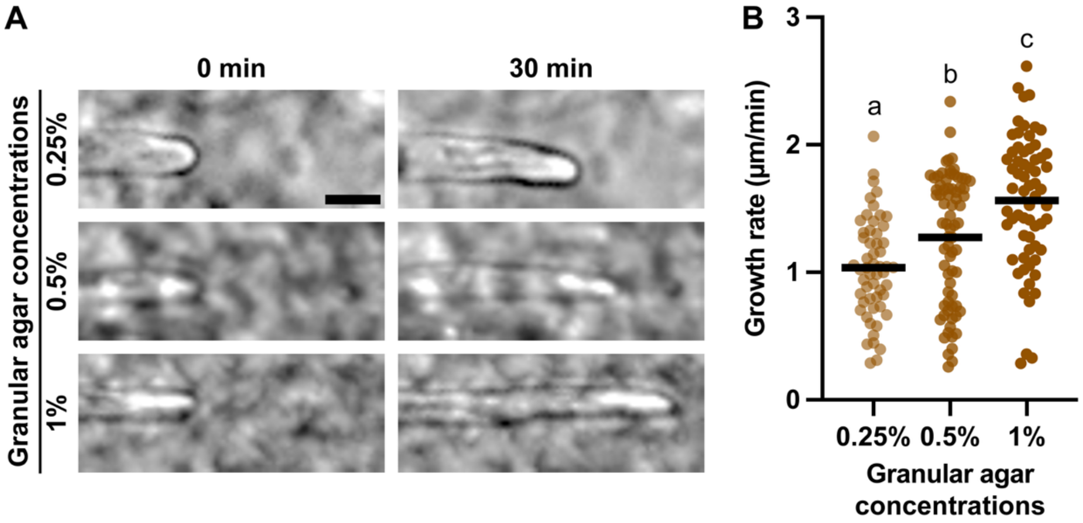


**Supplementary Fig. S4. Root hairs grow faster with the increased percentage of granular agar**

**A,** Representative brightfield images of root hairs growing in different granular media at two timepoints (Scale bar = 20 µm). **B,** Root hair growth rate quantified over 30 min. Each data point indicates one root hair. n = 55-73 root hairs from at least 10 seedlings from 4 independent experiments. Different letters denote significance, $p < 0.05$, ANOVA and post-hoc Tukey test.

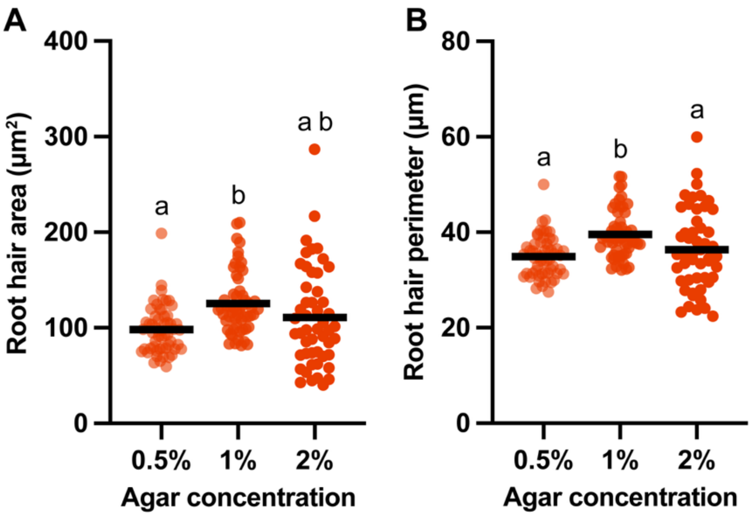


**Supplementary Fig. S5. Root hair cross-sectional area and perimeter show mixed trends with increasing stiffness of the growth environment**

**A,** Root hair cross-sectional area measured using XZ cross-sections of S4B stained, 5-day-old Col-0 roots. **B,** Root hair perimeter measured using the same method as for root hair cross-sectional area. Each data point represents one root hair. $n \geq 51$ root hairs from at least 15 seedlings from 5 independent experiments. Different letters denote significance, $p < 0.05$, ANOVA and post-hoc Tukey test.

**Supplementary Table S1:** Outcome of Relative Weight Analysis shows that turgor pressure is the major influencer to the surface stress in root hair FEM and shows a positive correlation.

| Variables | Raw.RelWeight | Rescaled. RelWeight | Sign | Sign.Rescaled. RelWeight |
|---|---|---|---|---|
| **Turgor pressure** | 0.48324616 | 49.427177 | + | 49.427177 |
| **Cell wall thickness** | 0.46233445 | 47.288294 | – | –47.288294 |
| **Circumferential modulus** | 0.03211261 | 3.284528 | – | – 3.284528 |

**Supplementary Table S2:** Individual values of turgor pressure (MPa) experimentally observed in trichoblasts and used to run root hair FEMs.

| | **Agar concentrations** | | |
|---|---|---|---|
| | **0.5%** | **1%** | **2%** |
| **Independent experiment - 1** | 0.80 | 0.76 | 0.64 |
| **Independent experiment - 2** | 0.91 | 0.77 | 0.75 |
| **Independent experiment - 3** | 0.87 | 0.79 | 0.62 |
| **Independent experiment - 4** | 0.89 | 0.78 | 0.65 |
| **Independent experiment - 5** | 0.83 | 0.77 | 0.58 |
| **Average** | 0.86 | 0.77 | 0.65 |
